## Supplementary Information for "Leveraging supervised learning for functionally-informed fine-mapping of cis-eQTLs identifies an additional 20,913 putative causal eQTLs"

**Supplementary Files legend**

Supplementary File 1. List of selected feature names and their importances in each tissue
Supplementary File 2. List of credible sets where the number of variants became significantly smaller after updating the PIP using EMS as a functional prior in whole blood
Supplementary File 3. List of newly identified potentially causal/non-causal eQTLs
Supplementary File 4. List of candidate genes identified in co-localization analysis
Supplementary File 5. Abbreviation of the assay and the tissue names used in the study
Supplementary File 6. Details of grid search for hyperparameter tuning in the training of whole blood EMS

**Supplementary methods**

Here, we describe the steps performed in the case of whole blood tissue. We followed the same steps for all the 49 tissues. EMS denotes the EMS in Whole Blood, unless otherwise noted.

Fine-mapping of GTEx variants

Fine-mapping of GTEx v8 variants was performed in Ulirsch et al[1] in the following steps:

1. Filter out the genes that do not have any variants with association p-value < $5 \cdot 10^{-8}$
2. Calculate the in-sample LD while adjusting for the covariates including the PEER factors.
3. Run SuSiE[2] and FINEMAP[3] with uniform prior and with default parameters

For more detail, see Ulirsch et al[1].

Annotating variant-gene pairs

In order to construct a predictor for causal eQTLs, we first defined the positive and negative labels. Specifically, we defined positive variant-gene pairs as the ones with PIP >0.9 for both of the two methods (SuSiE and FINEMAP), and negative ones as those with PIP<0.0001 for both methods. Since negatively labeled samples were large, we downsampled to 10,000 variant-gene pairs. Then we annotated the positive and negative variant-gene pairs with 6,121 features. Continuous features were turned to log scale after removing the sign with $log_2(abs(x) + 1)$transformation. We did not include conservation related features, gene expression, or gene constraint in the feature set. These steps resulted in a binary classification problem to classify putative causal vs non-causal variant-gene pairs given 6,121 features. In other words, letting the number of features m=6,121, the number of variant-gene pairs be $n$, the binary label vector of size $1 \times n$ be $y$, and the feature matrix of size $n \times m$ be $X$, we want the best $f(X)$ that maximizes the prediction accuracy of $y$.

We note that the PIP threshold for defining the positive and negative label is manually decided, considering the balance of the quality of the label and the number of positively labeled samples. Also, although PIP is a continuous measure that reflects the probability of being causal, we did not set the problem to be a regression problem to directly predict the PIP, because of the potential artifact of PIP mis-calibration and the highly skewed distribution of the PIP itself.

Feature selection for the predictor

We performed feature selection because of the high correlations among the features where a number of features are thought to be useful in only specific cell types (**Fig. S4**), and to reduce the complexity of the calculations. We performed feature selection in the following steps:

1. For baseline binary features, we excluded the histone mark-related features, TSS related features (as they overlap with distance to TSS or roadmap related features), as well as the conservation related features. (These features are not included in the count of the number of features.)
2. For each of the cell type-specific ROADMAP features, we calculated the precision, recall and F1 measure of the binary classification to characterize the enrichment of the feature in putative causal variants. Then, we selected variants in top 2.5% of the F1 measure or the precision as useful enriched features. We manually chose this and other thresholds based on the distributions of the statistics, leaving more features than what the elbow plot suggests (**Fig. S16**). To rescue a small number of features that are depleted in putative causal variants (and thus are also useful in prediction), we also selected the variants with odds ratio <0.5 and Fisher's exact test p-value < 0.05.
3. For each of the Basenji features, we first computed the feature importances, defined as the mean decrease of impurity (MDI)[4] of the binary prediction in a simple random forest model using the 5,313 Basenji features plus distance to TSS. The parameters for the simple random forest model in this step were set to 100 trees and 50% features per split (we did not optimize this forest, since it is purely for feature selection). We ranked all the Basenji features by the feature importance, and selected the top 100 features (since important features with high correlation to other features can have a decreased feature importances[4], we set the cut-off value to be larger than what elbow plot suggests).

This feature selection resulted in 152 features in Whole Blood and 149-157 features for other tissues (**Supplementary File 1**). We next tried to maximize the prediction accuracy of this binary classification problem by training the predictor, as described in the next section.

Training the predictor

We set the objective function to maximize in the prediction task to be the AUROC of the binary prediction (specifically, the average AUROC over 10 repeats of splitting the data into 70% training and 30% test, where negative samples were randomly sampled each time so that the ratio is 1:1, to reduce the computational burden). We tuned the hyperparameter of the random forest classifier to maximize the objective function, as described below:

1. Perform 100 iterations of random search, selecting hyperparameters randomly from the pre-defined parameter space;
   - number of trees: $N \in \{10, 32, 100, 316, 1000\}$ (corresponding to a log-linear increment)
   - minimum number of samples per split: $s \in \{2,8,16\}$
   - fraction of features to consider: $f \in \{0.1, 0.32, 0.64, 1\}$
   - maximum depths of each tree: $d \in \{2, \infty\}$
   - purity criteria $C \in \{Gini, Entropy\}$
2. Select the best (maximizes the AUROC) hyper-parameter from step 1, and perform exhaustive grid search on the grid defined as a function of the best hyper-parameter as follows:
   Let the superscript $_{best}$ note the best parameter found in step 1.
   number of trees: $N \in \{0.5N_{best}, 0.6N_{best}, \ldots, 1.4N_{best}, 1.5N_{best}\}$
   minimum number of samples per split: $s \in \{max(2, 0.5s_{best}), 0.5s_{best} + 2, 0.5s_{best} + 4, \ldots, 2s_{best}\}$
   fraction of features to consider: $f \in \{0.5f_{best}, 0.6f_{best}, \ldots, f_{best}\}$,
   while fixing the two parameters:
   maximum depths of each tree: $d = d_{best}$
   purity criteria $C = C_{best}$
3. Accept the hyper-parameter pairs with the best performance (maximizes the AUROC) over all the parameter combinations tested.

The record of parameter tuning for EMS in whole blood tissue can be found in **Supplementary File 6**. The final predictor is trained with the best hyper-parameter set using 100% of the data. We note that using different metrics (e.g. area under precision-recall curve = AUPRC) might result in slight differences in the predictor error mode, and therefore applied the scaling process as described in the next section.

### Scaling the score

The random forest output reflects the estimated probability of a variant-gene pair being causal, but is not scaled properly due to the downweighting process described above. In order to let the score be interpreted as the probability of a variant-gene pair being putative causal, we fixed the random forest predictor architecture, repeated the process of dividing the data into 90% training and 10% test data for 100 times, and observed the distribution of the positive and negative labeled test data for each output score bin (we set the bin size to be 0.01 with a moving window of 0.001). We then defined EMS as the number of positive samples in the bin divided by the total number of samples in the bin, after re-scaling the negatives to cancel out the downsampling effect. In other words, let $C$ be the set of positively labeled pairs, and $x$ be a random draw of a variant-gene pair, $B$ be a score bin, then

$EMS(B) = prob(x \in C \mid x \in B)$.

When the total number of variants in GTEx dataset is $N$, the total number of positively labeled samples is $n_p$ and the negatively labeled samples were downsampled to fraction $r$, the number

of positively labeled samples in the bin is $n_p(B)$, and the total number of samples we observe in the downsampled dataset in the bin is $N(B)$, the $EMS$ can be computed as
$EMS(B) = prob(x \in C \mid x \in B) \approx \frac{n_p(B)}{n_p(B) + (N(B)-n_p(B)) \cdot r^{-1}}$, considering the fact that $n << N$.
Thus, EMS for a variant-gene pair in a tissue is the probability that a given variant in GTEx v8 dataset will have PIP>0.9 for the expression of the gene in the tissue in statistical fine-mapping by SuSiE and FINEMAP with uniform priors.

The distribution of the putative causal variants along with the different steps of the scaling processes described above is shown in **Fig. S17.** We interpolated the score when needed. Along with this EMS, we also provide normalized EMS, defined as EMS divided by $prob(x \in C) = \frac{n_p}{N}$, corresponding to the enrichment of EMS compared to the uniform prior belief. This normalized EMS was used in **Fig. S18** as a sanity check for easier interpretation.

Enrichment analysis in replication cohorts

Geuvadis fine-mapping data was downloaded from http://www-personal.umich.edu/~xwen/geuvadis/. We defined the variant-gene pairs with PIP>0.9 in their fine-mapping annotation described in the web page. Fine-mapping of BBJ traits is described in Kanai et. al[5]. We followed the steps as the UKBB data for the enrichment analysis; for each (non-coding) variant, we calculated the maximum PIP over all the hematopoietic traits, as well as the maximum Whole-Blood EMS over all the genes in the cis window of the variant, and defined a variant as putative hematopoietic trait-causal if it has SuSiE PIP higher than 0.9 in any of the hematopoietic traits. We focused on the variants that exist in the GTEx v8 dataset to reduce the calculation complexity. Finally, for both two cohorts, the variants other than putative causal ones were randomly downsampled to achieve a total number of variants to be exactly 100,000. We note that we did not perform the filtering for BBJ variants, because of the small number of putative causal variants (n = 64) intersected with the GTEx variants (which are predominantly from European samples).

Approximate functionally informed fine-mapping with EMS

Here we introduce our algorithm for approximate functionally-informed fine-mapping, adapted from the SuSiE method [2] and analogous to the BFMAP method [6]. We then explain how we used this method to perform approximate functionally informed fine-mapping of GTEx data using EMS as a prior.

The SuSiE model is introduced in ref [2]; we briefly review it here. Let $y$be the $n \times 1$ phenotype vector and $X$be the $n \times m$ genotype  matrix, where $n$ and $m$ are the number of individuals and the number of variants in the locus, respectively. Let $\beta$ be the vector of length $m$ of variant effects and $\epsilon$ be a noise vector of length $n$. Then the SuSiE model can be written as:

$$y = X\beta + \epsilon$$
$$\epsilon \sim N(0, \sigma^2 I_n)$$

$$\beta = \sum_{l=1}^{L} \beta_l$$
$$\beta_l = \gamma_l b_l$$
$$\gamma_l \sim Multinomial(1, \pi)$$
$$b_l \sim N(0, {\sigma^2}_{0,l})$$

Here, $\pi$ denotes the vector of prior probabilities of each variant being the causal variant for a single effect. The uniform prior, $\pi = (\frac{1}{m}, \frac{1}{m}, \ldots, \frac{1}{m})$, is the default setting.

The SuSiE algorithm computes a variational approximation in which the posteriors for the $\beta_l$ are independent of each other. Under this approximation, let

$$\alpha_l{}^{\pi}(v) = prob(\gamma_l = e_v | D; \pi)$$

where $e_v$ is the $v$-th standard basis vector and $D$ denotes the data. (The superscript $\pi$reflects the prior, and does not denote the power. For simplicity, we omit the dependence on $\sigma^2$ and ${\sigma^2}_{0,l}$.)

A credible set $C_l$ is computed for each $l$ from the $\alpha_l{}^{\pi}$, by iteratively adding the variant corresponding to the largest remaining element of $\alpha_l{}^{\pi}$ until the sum exceeds the pre-defined threshold, typically set to 0.95. The PIPs are computed from the $\alpha_l{}^{\pi}$:

$$PIP_{\pi(v)} = prob(\beta_v \neq 0 \mid D; \pi)$$
$$= 1 - \prod_{l=1}^{L} (1 - \alpha_l{}^{\pi}(v))$$

The SuSiE model fixes the number of effects (which approximately equals the number of causal variants in the locus) at $L$ for inference and then does a post-hoc pruning step to determine which $\alpha_l$most likely do not correspond to true effects. Specifically when $\alpha_l$ corresponds to a true effect, then most of the mass of $\alpha_l$ is concentrated around variants in high LD with each other; conversely, if $\alpha_l$ does not correspond to a true effect then the weights $\alpha_l(v)$ are typically spread across many variants. Thus, a credible set is called “pure” if all pairs of variants in the credible set are correlated at $R^2 > 0.25$; otherwise, the credible set is called “impure”. If $C_l$ is impure, then we conclude that $\alpha_l$ does not correspond to a true effect.

Let $u = (\frac{1}{m}, \frac{1}{m}, \ldots, \frac{1}{m})$, and let $f$ denote an $m$-vector of functionally-informed prior probabilities. Here, we will show that if we have already computed $\alpha_1{}^u, \ldots, \alpha_L{}^u$, then we can quickly approximate $\alpha_1{}^f, \ldots, \alpha_L{}^f$, which we can then use to compute functionally-informed PIPs and credible sets (the superscripts $u$ and $f$ denotes the uniform and functional prior, not the power).

To do this, we write:

$$\alpha_l{}^{\pi}(v) = prob(\gamma_l = e_v \mid D \,;\, \pi)$$
$$\propto prob(D \mid \gamma_l = e_v \,;\, \pi) \cdot prob(\gamma_l = e_v \,;\, \pi)$$
$$= prob(D \mid \gamma_l = e_v) \cdot \pi(v)$$

where the normalization is such that the vector $\alpha_l{}^{\pi}$ sums to one. Thus,

$$\alpha_l{}^{u}(v) \propto prob(D \mid \gamma_l = e_v)$$

Which allows us to conclude that

$$\alpha_l{}^f(v) \propto \alpha_l{}^u(v) \cdot f(v)$$

i.e.,

$$\alpha_l{}^f(v) = \frac{\alpha_l{}^u(v) \cdot f(v)}{\sum_{j=1}^{m} \alpha_l{}^u(j) \cdot f(j)}$$

If the variational approximation given by SuSiE is exact, then this will be an exact computation. If it is not exact, then the results of this procedure will not necessarily be the same as if SuSiE had been applied with $\pi = f$.

We use $\alpha_l{}^f$ to compute PIPs and credible sets as in the original SuSiE algorithm, with one change. Under a uniform prior, if $\alpha_l$ does not correspond to a true effect, then its entries are typically spread over many variants, with no variant getting too much weight. Since loci typically have many variants, such $\alpha_l$ do not have much impact on the PIP. However with a functionally-informed prior, $\alpha_l$ can have large entries for variants given a high prior value, even if $\alpha_l$ does not correspond to a true causal effect.

Thus, we set
$\hat{\alpha}_l{}^f = \frac{\alpha_l{}^u(v) \cdot f(v)}{\sum_{j=1}^{m} \alpha_l{}^u(j) \cdot f(j)}$ if $\alpha_l{}^u$corresponds to a pure credible set, and
$\hat{\alpha}_l{}^f = \alpha_l{}^u$ otherwise.
We then use $\hat{\alpha}_l{}^f$ to compute PIPs and credible sets.

Finally, since EMS has a unit of estimated probability of being causal, it can be directly plugged in as the functional prior (we set the $L = 10$ as in the SuSiE default setting).

Previous work[7] in functionally informed fine-mapping adjusted the prior so that the maximum prior value did not exceed 100 times the minimum prior value. We also conducted a second round of functionally informed fine-mapping with a similar adjustment of the prior, for comparison; **Fig. S8**. To obtain a max/min ratio of $R$, we simply scaled the functionally-informed prior $f$ with the constant power. i.e. let the largest prior across genome be $\Phi_G$and the smallest be $\phi_g$, and the updated weight be $\hat{f}(1)$,..., $\hat{f}(m)$, then
$\hat{f}(v) = f(v)^c$, where $c = log_{\frac{\Phi_G}{\phi_g}} R$.

Limitation and possible next steps for the work

In this section we will discuss the details and possible next steps regarding to the limitations of the study:

1. The GTEx cohort is highly biased towards aged adult samples, which makes it hard to generalize the EMS across different developmental stages and ages. Also, the power to call a putative causal variant in a statistical fine-mapping is higher when the recombination rate and the allele frequency of the variant is high (we observed difference of EMS distribution in different allele

count in the GTEx sample; **Fig. S19**). Future work includes testing the generalizability of EMS in a cohort with not only distinct ethnicity but also age distribution, and also cohorts with larger sample sizes (that provides eQTLs with smaller allele frequency).

2. We envision that adding the effect size as an output of the predictor can be potentially applied for quantitative predictions of gene expression changes[8]. We also note that we have not tested all possible predictor architectures, nor all the parameter tuning methods, although random forest using full 6,121 features outperforms a number of major predictors (**Fig. S20**). Using different predictors and parameter-tuning methods could result in higher prediction power. Also we did not repeat the process of training the predictor and performing functionally informed fine-mapping considering the fact that the sample size is not likely to be the determinant of the prediction power (**Fig. S10**) and also to avoid potential mis-calibration of the PIP, such an approach with careful consideration might be useful in identifying a larger set of putative causal eQTLs.

3. Lastly, the process of building EMS and $PIP_{EMS}$ is independent from one tissue to another, although we observe similarity in the enrichment pattern of independent set of reporter assay-positive variants, eQTL and complex trait signals in biologically related tissues (**Fig. S6, S11, S21**). As previous eQTL studies[9,10] have utilized the correlated structure of the causal mechanisms between similar tissues, future work taking advantage of tissue correlations would be valuable.

**Supplementary Figures:**

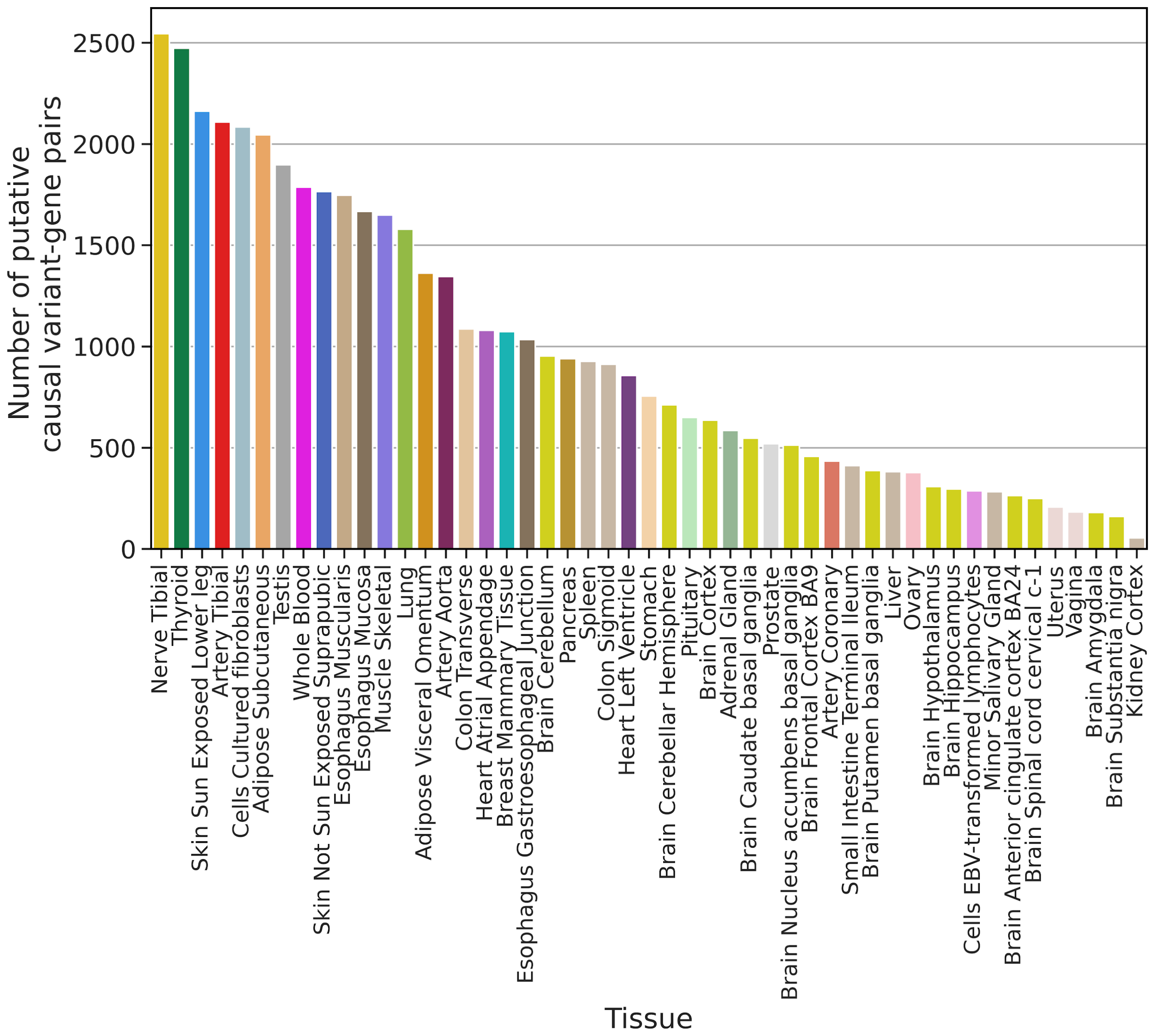


**Figure S1. Overview of fine-mapped eQTLs**
The bar chart shows the number of putative causal variant-gene pairs in different tissue, sorted in descending order.

a

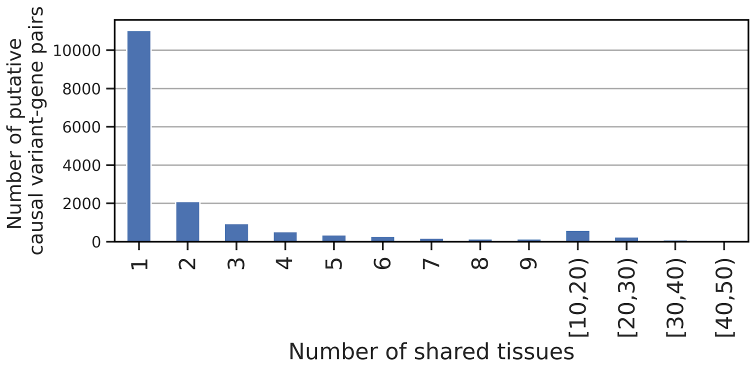


b

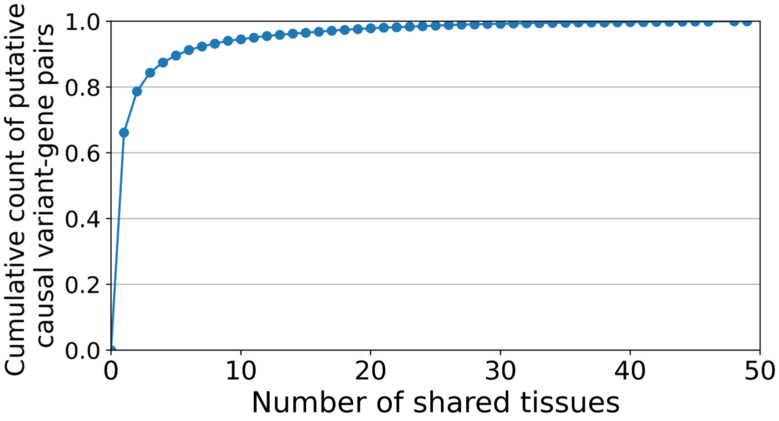


c

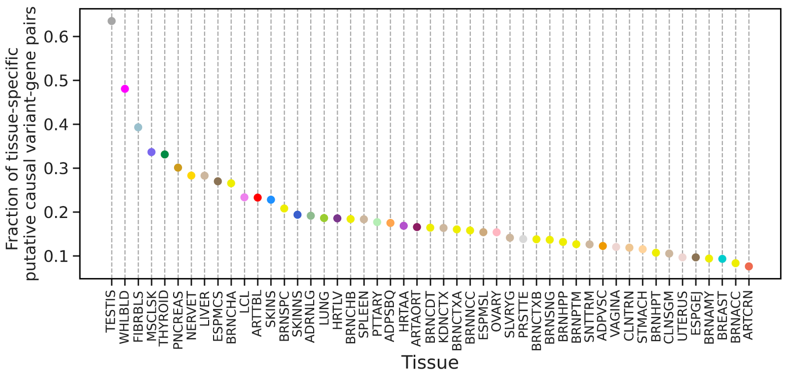


**Figure S2. Tissue specificity of the putative causal variants-gene pairs in GTEx v8**
**a. b.** The distribution (**a**) and the cumulative count (**b**) of the number of shared tissues for putative causal variant-gene pairs in GTEx . **c.** The fraction of tissue-specific putative causal variant-gene pairs for each tissue, sorted in ascending order. For panel **c** and all the other supplementary figures, the tissue name abbreviations can be found at **Supplementary File 5**.

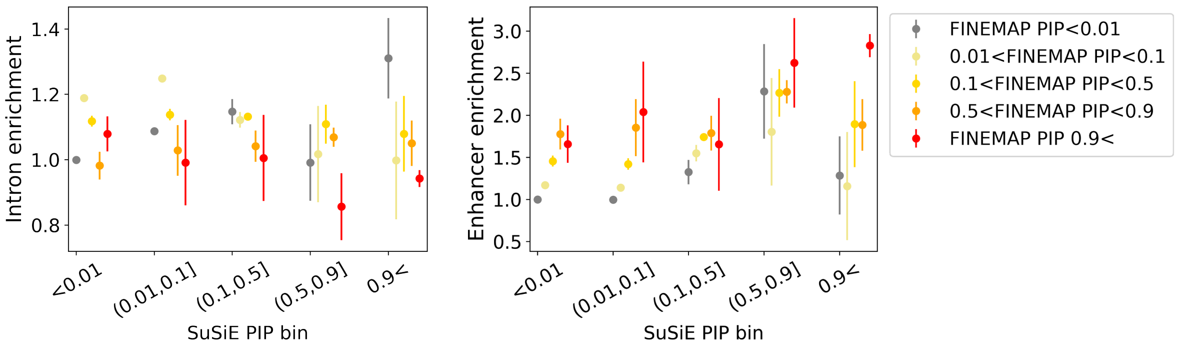


**Figure S3. Enrichment of functional annotations for variant-gene pairs with PIP bin of two different fine-mapping methods**

Enrichment of intron (left panel) or enhancer (right panel) features in variant-gene pairs with different PIP bins defined by SuSiE (x axis) and FINEMAP (color gradient) algorithms. In the bins with highest SuSiE PIP, the bin with lowest FINEMAP PIP has the highest enrichment of intron, and the bin with high FINEMAP PIP has the highest enrichment of enhancer, suggesting that taking the intersection of SuSiE and FINEMAP improves the accuracy of fine-mapping.

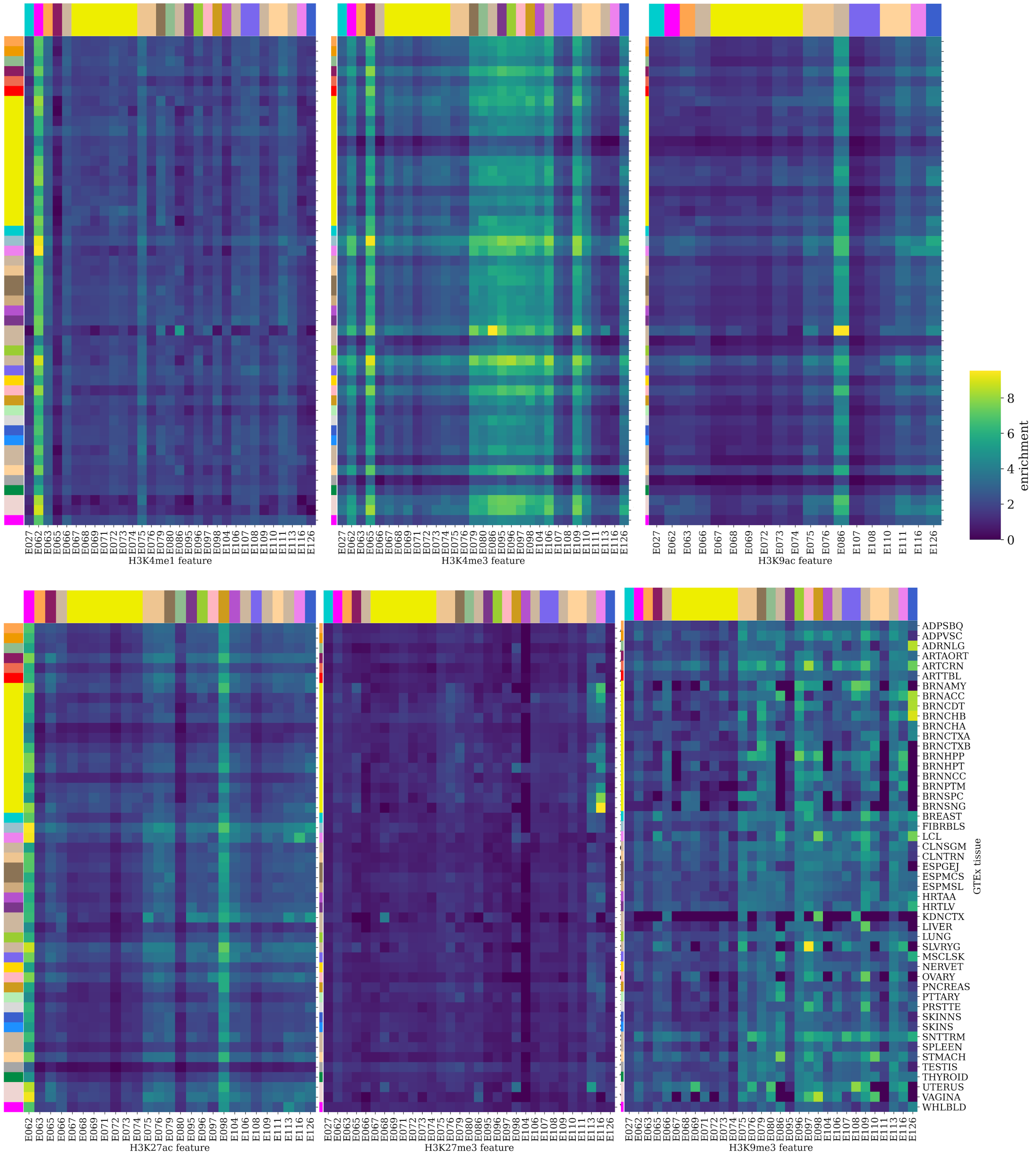


**Figure S4. Full overview of the tissue specific histone mark enrichment**
row = GTEx tissue, colored as in **Supplementary File 5**, column=tissue specific binary histone mark features from Roadmap, and the heatmap color shows the enrichment level of variant gene pairs with PIP>0.9 in each binary feature. See **Supplementary File 5** for the four letter abbreviation of the sample type.

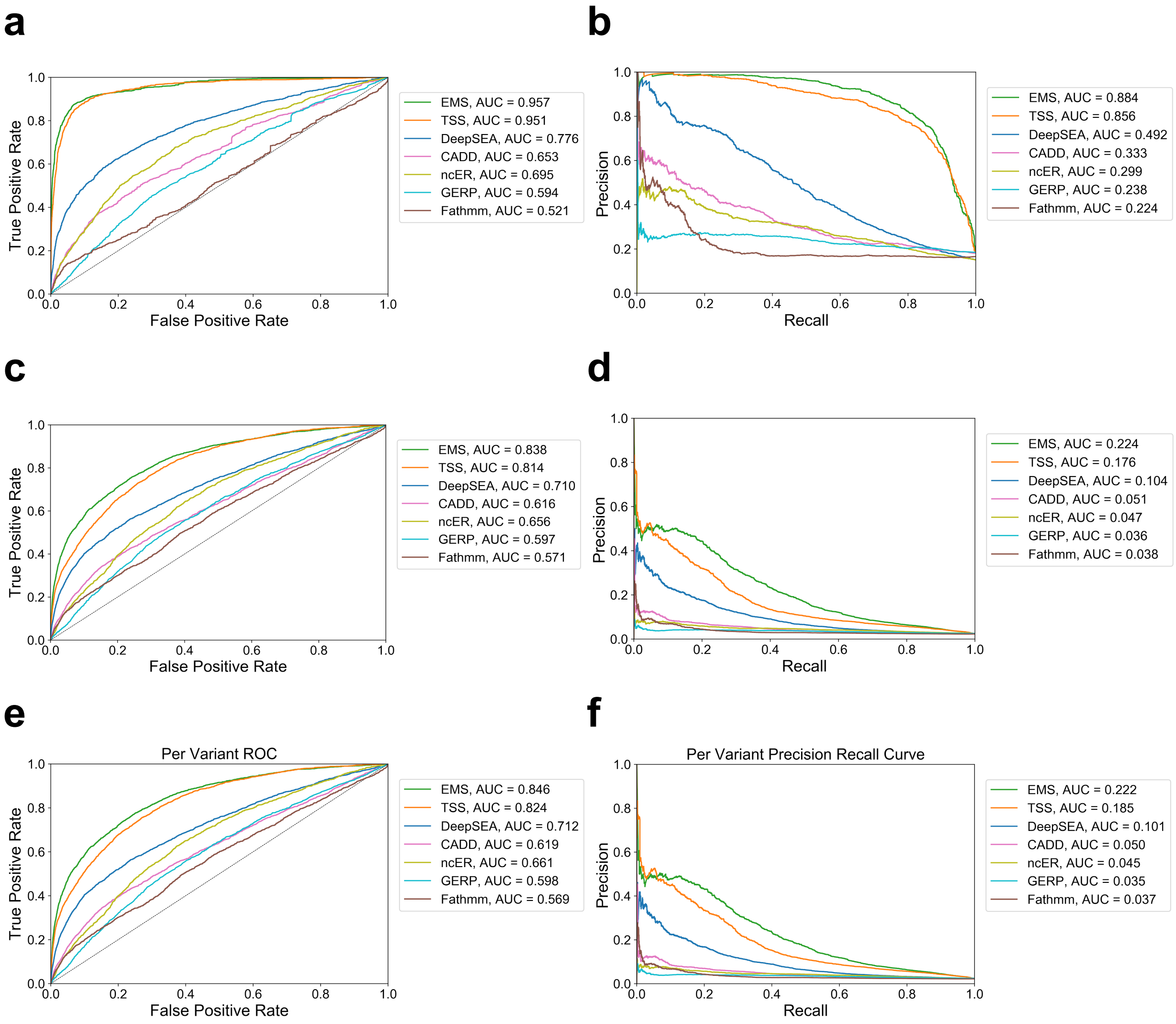


**Figure S5. Performance evaluation of EMS and other methods**

**a. b.** Receiver-operator characteristics (ROC) curve **(a)** and precision recall curve (PRC) **(b)** of the binary prediction of putative causal Whole-Blood eQTLs in GTEx v8 using different genomics scores including EMS. Here, negative variant-gene pairs are randomly down-sampled to n=10,000 (5.60x more negative variant-gene pairs than positive variant-gene pairs).

**c. d.** ROC **(c)** and PRC **(d)** of the same prediction problem as **a** and **b**, with increased number of negative samples. Here, negative variant-gene pairs are randomly down-sampled to n=998,213 (560x more negative variant-gene pairs than positive variant-gene pairs.).

**e. f.** AUROC **(e)** and AUPRC **(f)** of the binary classification of putative causal whole blood eQTLs variants in GTEx v8. Instead of letting each entry be a variant-gene pair, here we calculated the maximum EMS and minimum distance to TSS to represent the scores for each variant, and let each entry be a variant. (Whole Blood) EMS still shows the best performance, showing that EMS is useful not only for variant-gene pair prioritization but also for variant prioritization.

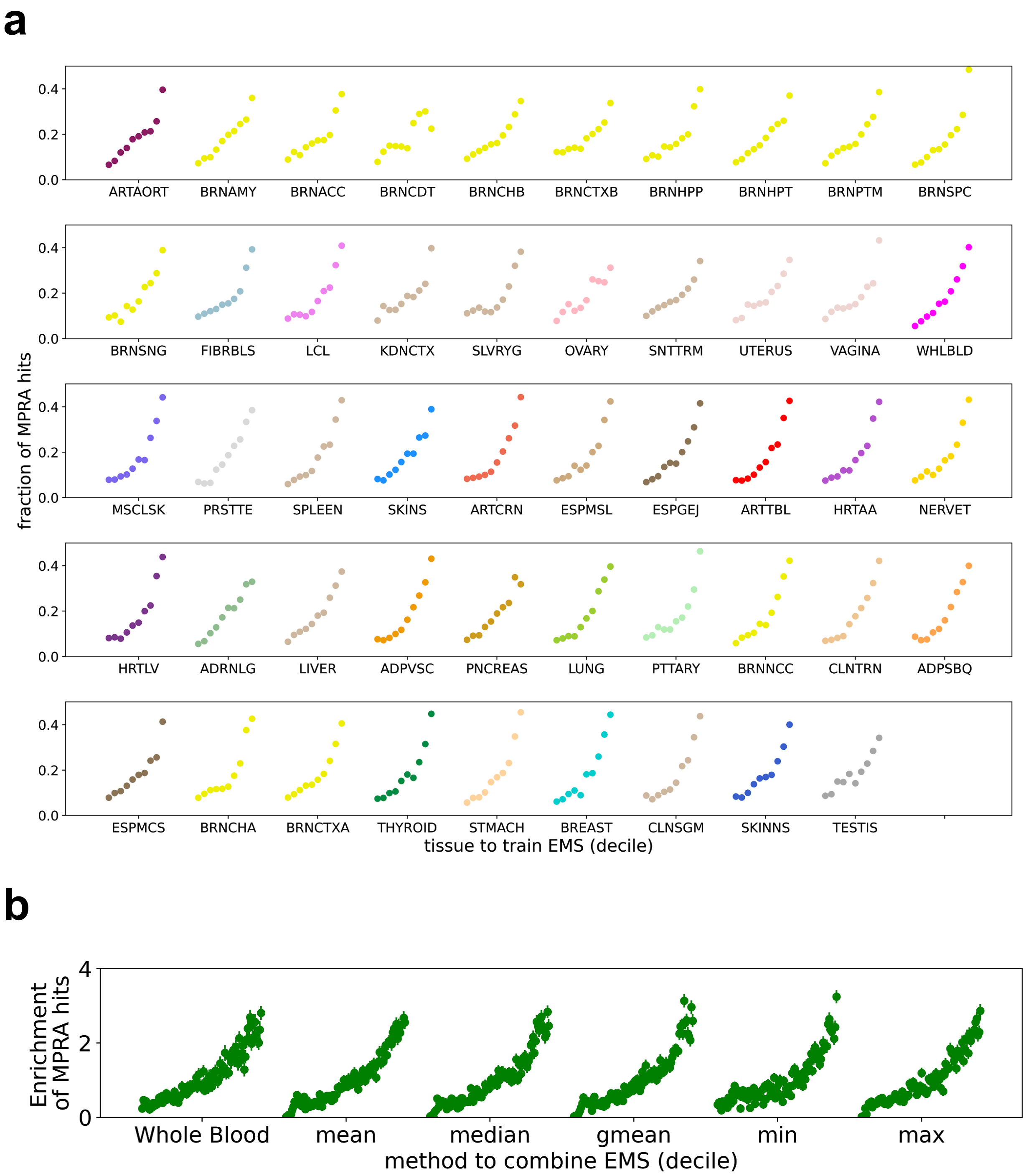
a
0.0
0.2
0.4
ARTAORT
BRNAMY
BRNACC
BRNCDT
BRNCHB
BRNCTXB
BRNHPP
BRNHPT
BRNPTM
BRNSPC
BRNSNG
FIBRBLS
LCL
KDNCTX
SLVRYG
OVARY
SNTTRM
UTERUS
VAGINA
WHLBLD
MSCLSK
PRSTTE
SPLEEN
SKINS
ARTCRN
ESPMSL
ESPGEJ
ARTTBL
HRTAA
NERVET
HRTLV
ADRNLG
LIVER
ADPVSC
PNCREAS
LUNG
PTTARY
BRNNCC
CLNTRN
ADPSBQ
ESPMCS
BRNCHA
BRNCTXA
THYROID
STMACH
BREAST
CLNSGM
SKINNS
TESTIS
fraction of MPRA hits
tissue to train EMS (decile)
b
0
2
4
Enrichment of MPRA hits
Whole Blood
mean
median
gmean
min
max
method to combine EMS (decile)

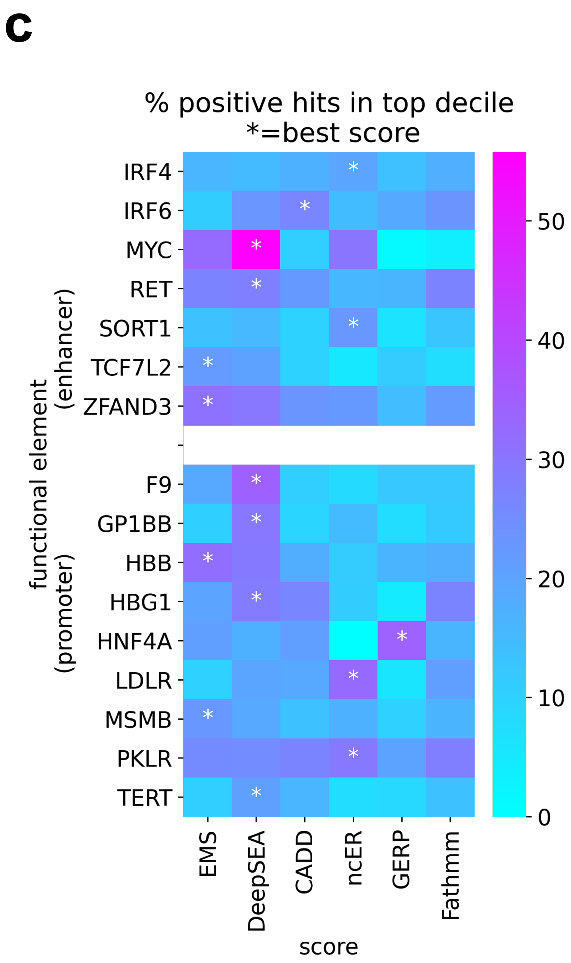


**Figure S6. Performance evaluation of EMS and other methods in prioritization of regulatory variants identified in saturation mutagenesis experiment**

**a. b.** Comparison of the performance of Whole Blood EMS against EMS trained on different tissues (**a**), and its mean, median, geometric mean, minimum and maximum across tissues (**b**) in prioritizing regulatory variants from publicly available targeted massive parallel reporter assay (MPRA) saturation mutagenesis results. **c.** Comparison of the different scoring methods, for each targeted functional element (EMS: geometric mean across 49 tissues, which empirically worked best in **b**). Column shows the different functional elements, and color shows the percentage of MPRA hits in the top score decile (within each element). Functional elements with <50 MPRA hits were removed in the plot.

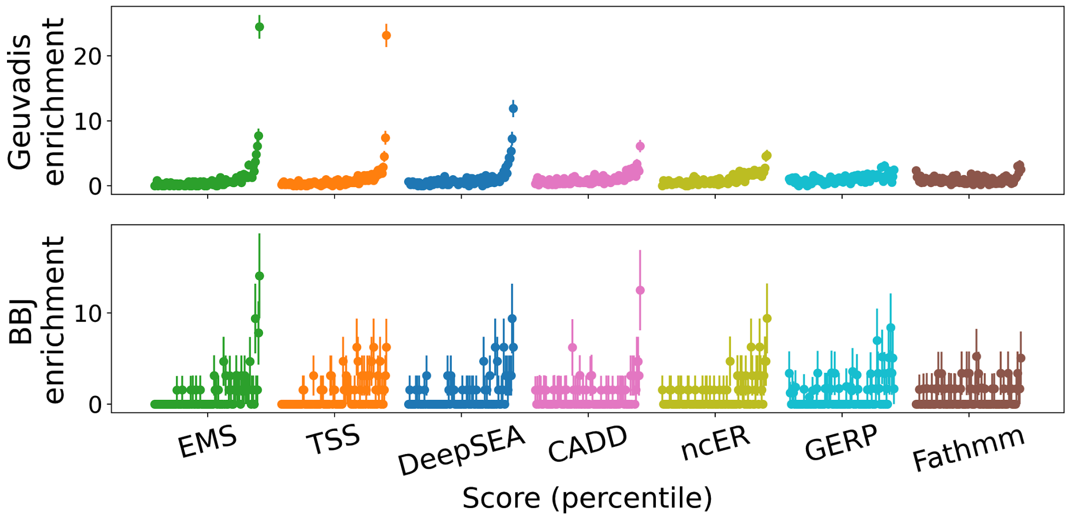


**Figure S7. Replication of performance comparison of EMS in putative causal regulatory variant prioritization versus putative causal complex-trait variant prioritization in a different cohort**

Enrichment of different scores (EMS, distance to TSS, DeepSEA, Fathmm, CADD, ncER and GERP) in putative lymphoblastoid cell line (LCL) eQTLs in Geuvadis (upper panel), and putative hematopoietic trait causal variants in Biobank Japan (BBJ; lower panel) dataset are shown.

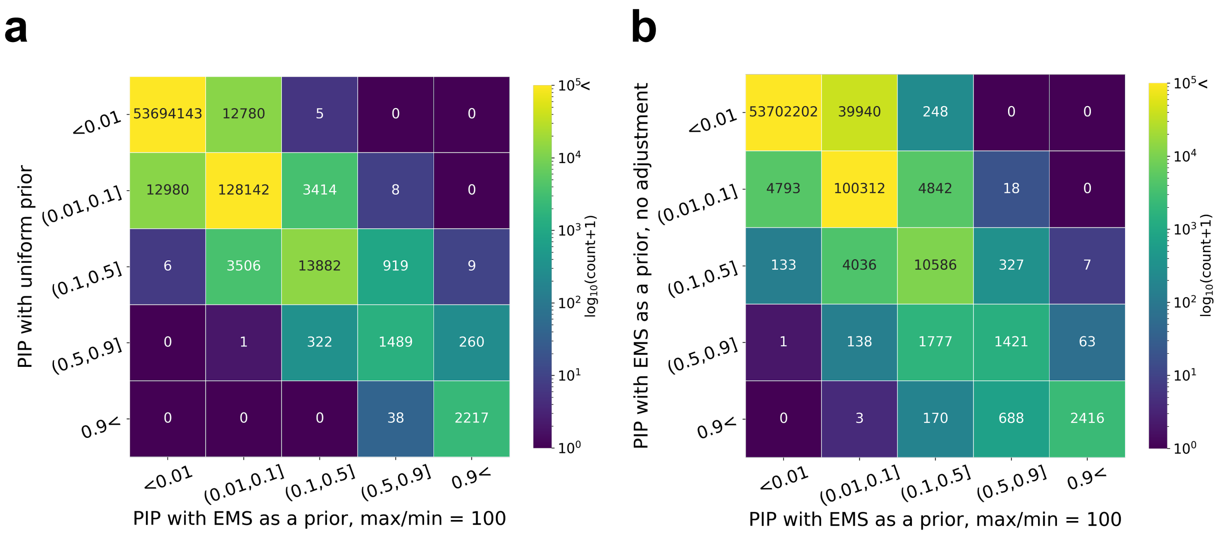


**Figure S8. functionally informed PIP with conservative max/min ratio in the prior**

**a.** Number of variant-gene pairs in different PIP bins (row = PIP with uniform prior, column = PIP with functionally informed but conservative prior, scaling the raw EMS so that the max/min ratio does not exceed 100).

**b**. Number of variant-gene pairs in different PIP bins using two different max/min ratio of the prior (row = PIP with raw EMS as a prior, column = PIP with conservative prior, identical to the column in **a.**). The max/min ratio was ~=40,000 in the raw EMS.

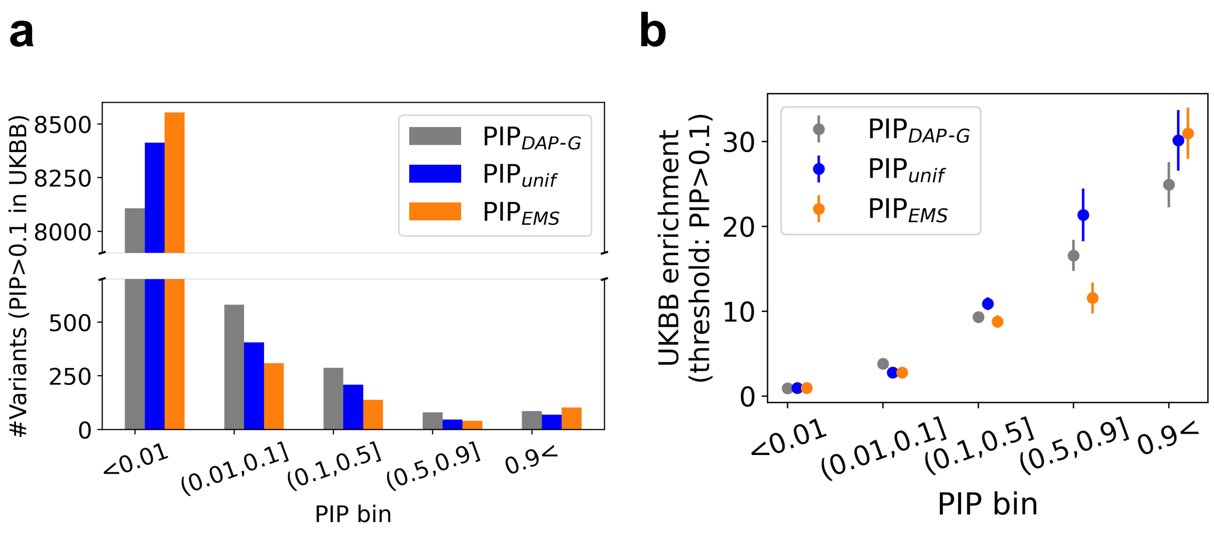


**Figure S9. Evaluation of $PIP_{EMS}$ using complex trait-causal variants in UKBB**

**a.** Number of variants with hematopoietic trait PIP>0.1 in UKBB, for different PIP bins (grey: publically available eQTL PIP using DAP-G, blue: PIP with uniform prior, orange: PIP with EMS as a prior). y axis is truncated.

**b.** Enrichment of variants with hematopoietic trait PIP>0.1 in UKBB, for different PIP bins (color: same as **a**).

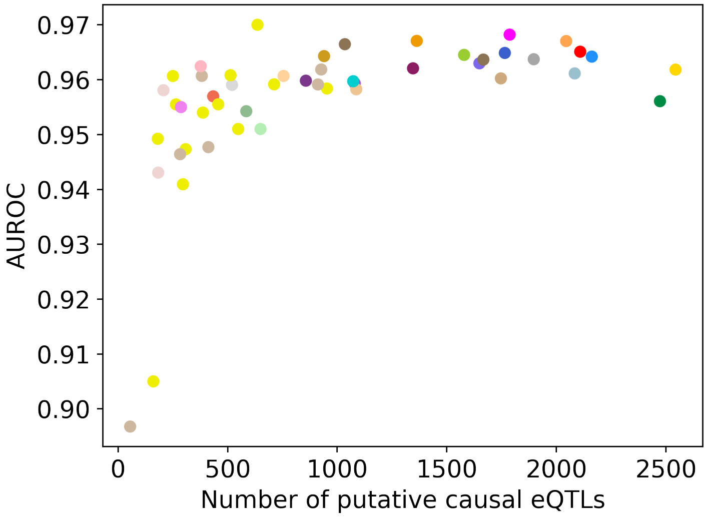


**Figure S10. Relationship between the positive training data size and the prediction accuracy of EMS in different tissues**

x axis shows the number of putative causal eQTLs, y axis shows the AUROC of the prediction in the held-out dataset, and different colors correspond to different tissues, as in **Fig. 5**. The average over 10 fold performance evaluation with 70% training and 30% test data split is shown. The overall correlation is weak (pearson r = 0.495, Fisher's exact test p=$2.99 \cdot 10^{-4}$), and is not significantly different from zero when filtering out the tissues with n<1000 (19 tissues, pearson r=$1.69 \cdot 10^{-4}$, p>0.05), suggesting that the positive training sample size is not the major determinant of the prediction accuracy, once it reaches a sufficient threshold.

**a**

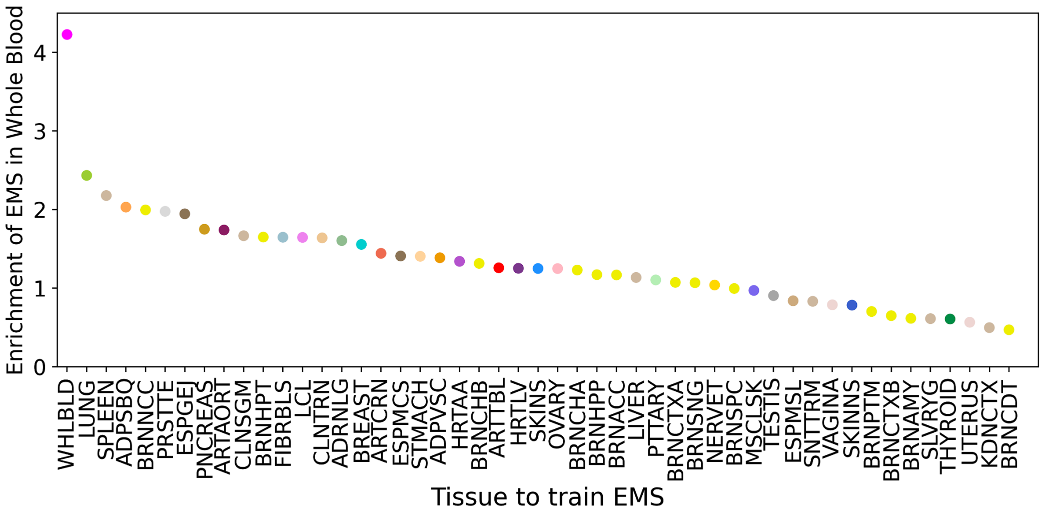


**b**

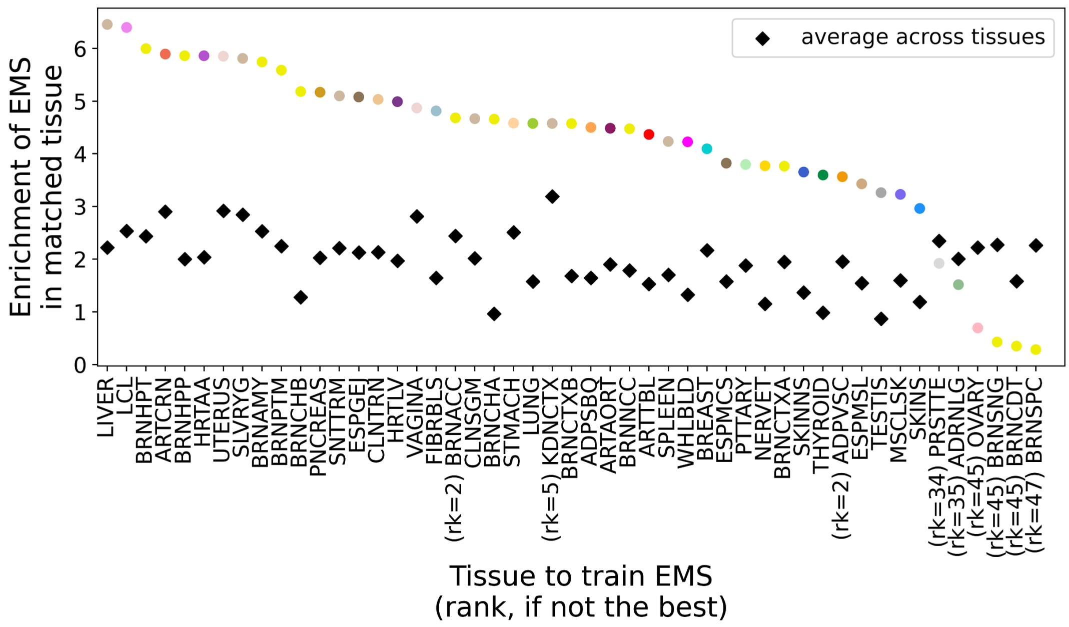


**Figure S11. Tissue specificity of EMS**

**a.** Enrichment of EMS that are trained on different tissues in the putative causal eQTLs in whole blood. Here, enrichment is defined as the log ratio of the median EMS in putative causal variants in whole blood and the median EMS in putative non-causal variants in whole blood. **b.** Enrichment of EMS that are trained on different tissues in the putative causal eQTLs in the same corresponding tissue. The median enrichment score across 49 tissues are shown in black, and the rank of the corresponding tissue is shown, when it does not show the highest enrichment.

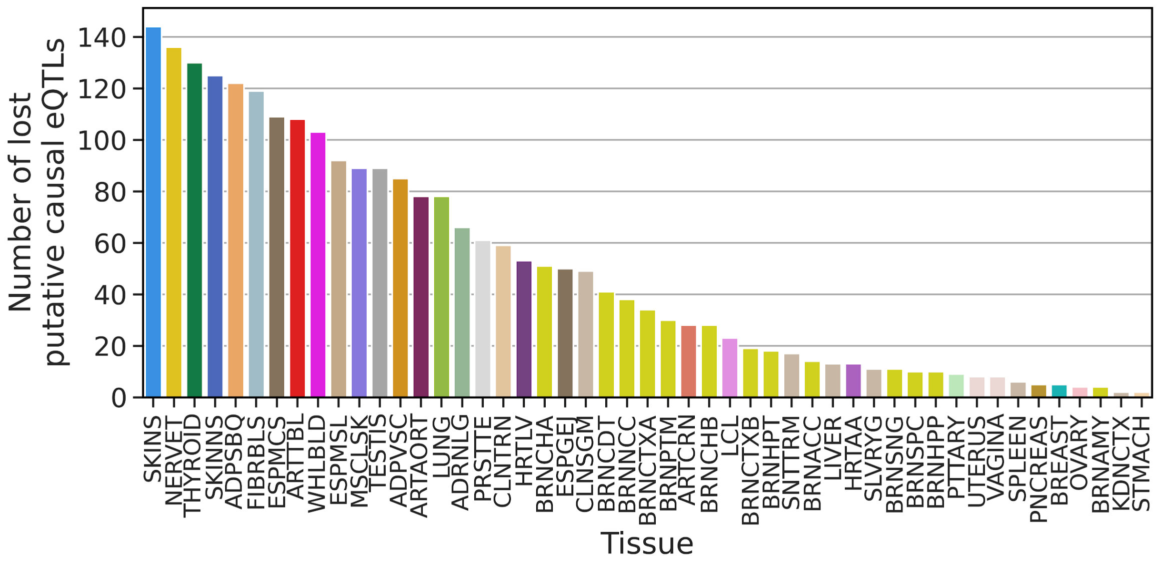


**Figure S12. Number of newly discovered "lost" putative causal eQTLs using $PIP_{EMS}$ per tissue**

x axis shows the tissue, and y axis shows the number of newly discovered "lost" putative causal eQTLs (**b**; $PIP_{EMS}$<0.9 and $PIP_{unif}$>0.9).

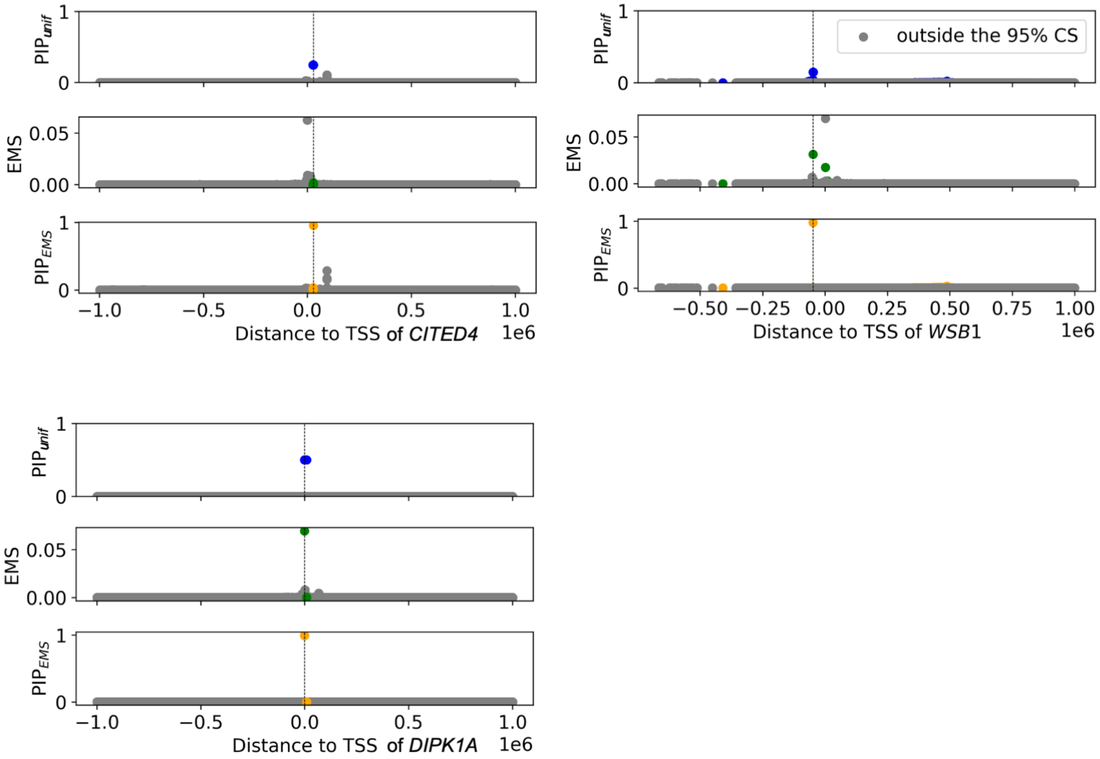


**Figure S13. +-1Mb window around the TSS for the example genes with newly identified putative causal eQTLs**

X axis is the distance to the TSS, and y is the PIP with uniform prior (top), EMS (middle), and PIP with EMS as a prior (bottom). Variants outside of the 95% credible set were greyed out.

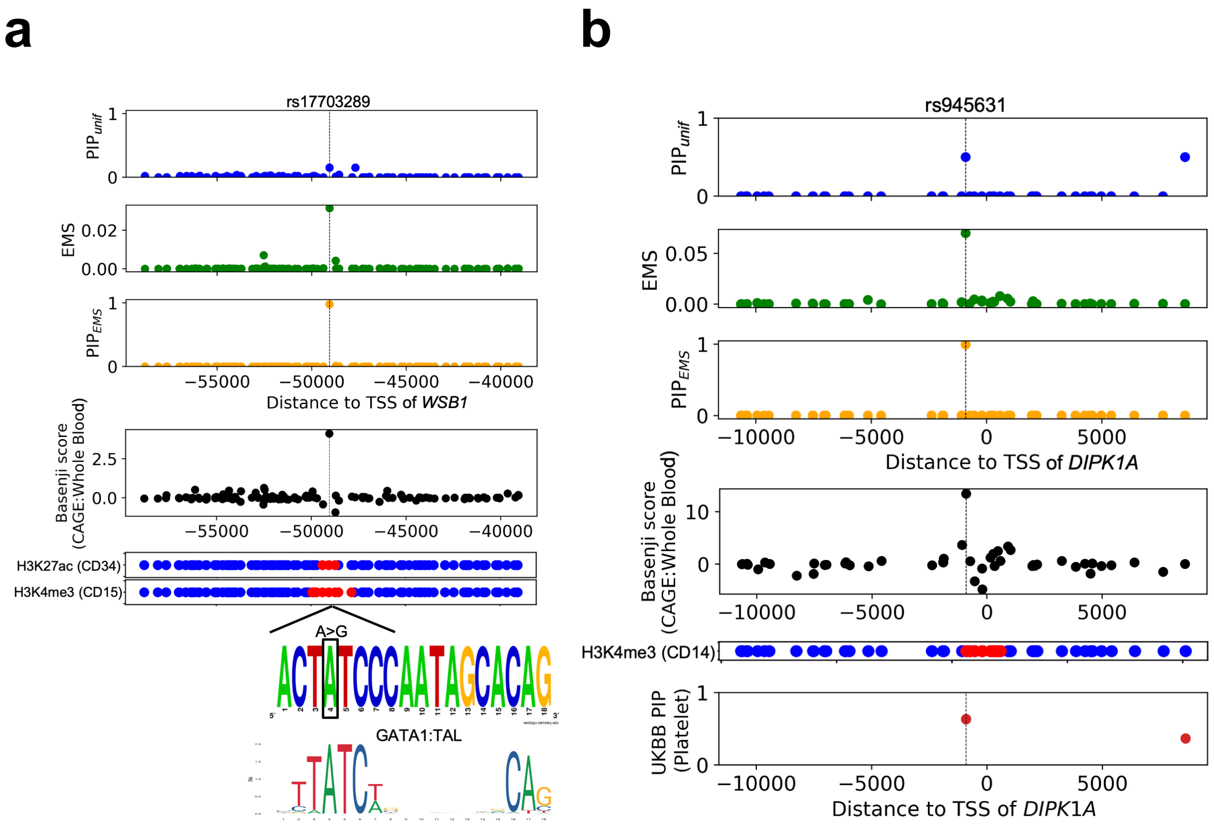


**Figure S14 Additional examples of putative causal eQTL prioritized by EMS**

An example of an upstream variant (rs17703289) of *WSB1* gene (left), and an intron variant (rs945631) of *DIPK1A* gene (right) showing high $PIP_{EMS}$ (>0.99). GATA1:TAL is known to be associated with hematopoietic processis[11], and rs945631 is associated with platelet count in UKBB (p=$9.8 \cdot 10^{-14}$, $PIP_{unif}= 0.64$, the highest in the credible set), each suggesting that the variant prioritized by $PIP_{EMS}$ is likely to have functional activities related to whole blood.

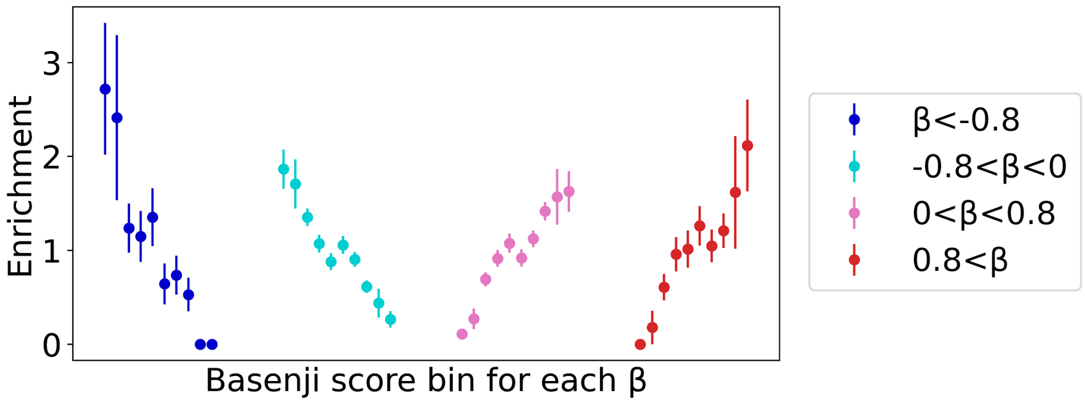


**Figure S15. Basenji score is predictive for the effect size (β) and direction of putative causal eQTLs**

For each (marginal) effect size bin of the putative causal eQTLs denoted in different colors, x axis is the basenji score bin for CAGE in neutrophil (highest: >20, lowest: <-20), and y axis is the enrichment of putative causal eQTLs in the effect size bin. Putative causal eQTLs with negative effect sizes (blue and light blue) are enriched in low basenji score bins, whereas those with positive effect sizes (pink and red) are enriched in high basenji score bins.

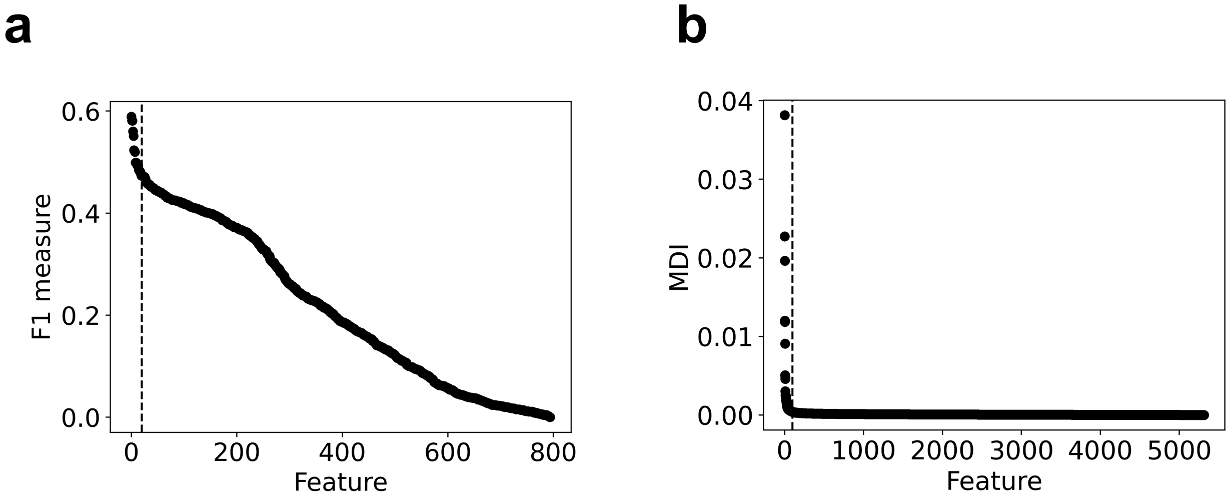


**Figure S16. Quality control in the feature selection step of EMS**

**a.** F1 measure of all the 795 ROADMAP features, sorted in descending order.

**b.** Feature importances (mean decrease of impurity = MDI) of all the 5,313 basenji features, sorted in descending order.

Dashed line shows the cutoff for accepting features.

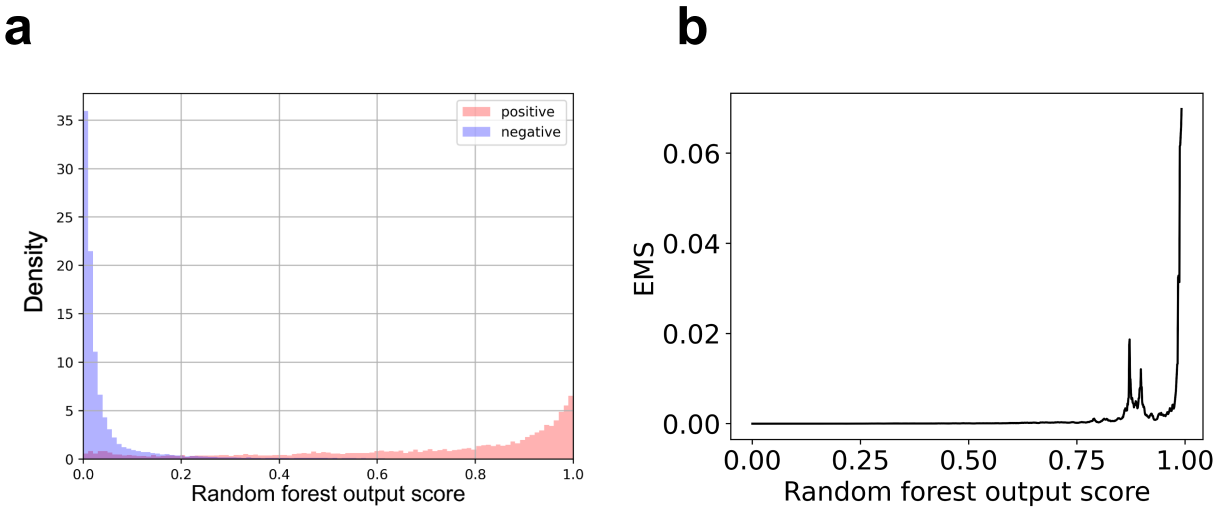


**Figure S17. Details of the scaling step of EMS**

**a.** The distribution (density) of negative (blue) vs positively (red) labeled data (100 repeats of 90% training and 10% test data split) as a function of raw random forest output score (x axis).

**b.** EMS as a function of raw random forest output score (x axis).

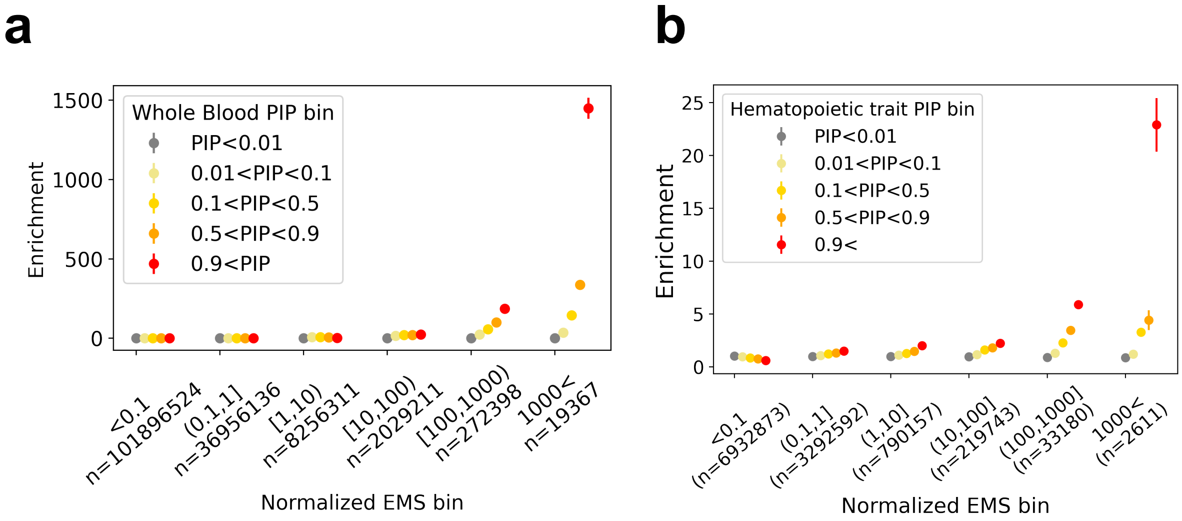


**Figure S18. Enrichment of eQTLs and hematopoietic trait-associated variants in EMS bins, by PIP bin**

The x axis is the normalized EMS bin (EMS divided by the probability of putative causal variant-gene pair in a random draw), and the y axis is the enrichment of variants in different PIP bin (**a** is for whole blood eQTLs in GTEx v8 and **b** is for hematopoietic trait in UKBB). The total number in **b** is smaller than that in **a**, since only the variants that exist in both datasets are included, and also the filtering to PIP>0.0001 was applied, as in Ulirsch et al[1].

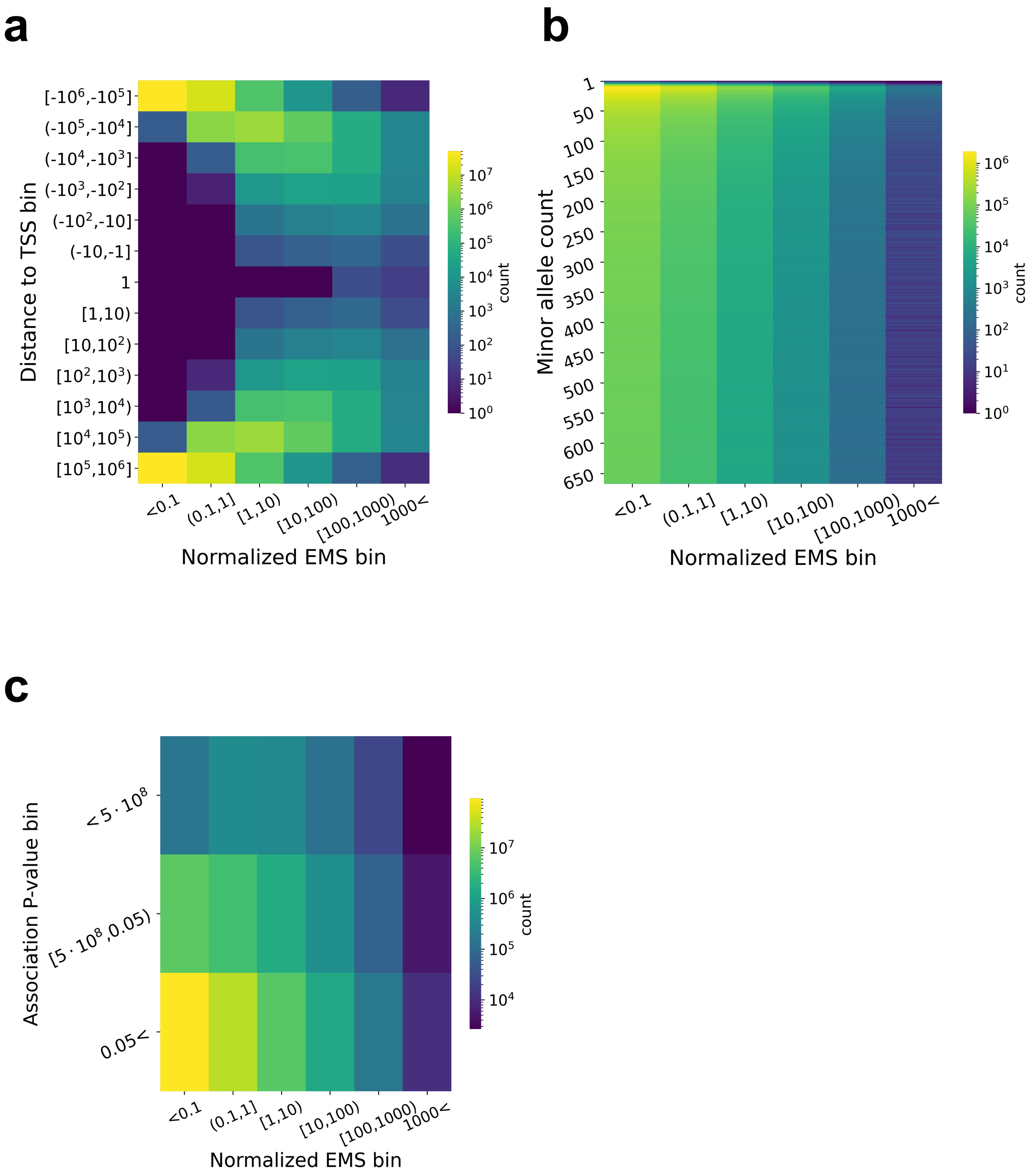


**Figure S19. Distribution of distance to TSS, allele count and association p-value of variant-gene pairs with different (normalized) EMS bin**

x axis is the normalized EMS bin, and y axis is the TSS distance (**a**), allele count (**b**) and association p-value (**c**) bin.

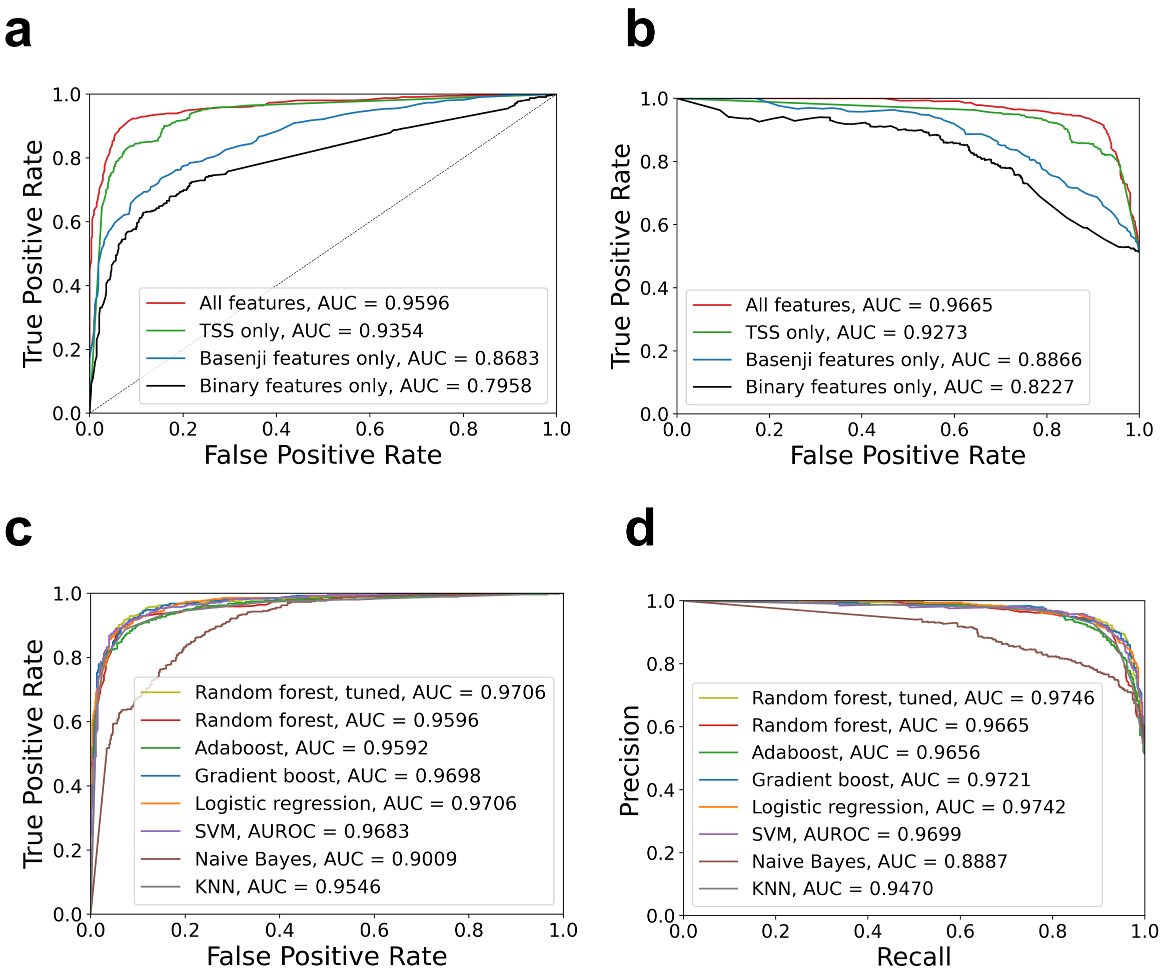


**Figure S20. Performance of Random forest compared in different feature sets and to different methods**

**a. b.** Performance comparison of the random forest classifier as a function of features to feed in, as ROC (**a**) or PRC (**b**). We fixed the random forest architecture (=default setting of scikit learn package v0.21.3), and compared the performance. Using all four types of feature categories (= EMS's approach) together resulted in the best performance.

**c. d.** Performance comparison of the random forest classifier against other major predictors, as ROC (**c**) or PRC (**d**). We fixed the classifier architecture (=default setting of scikit learn package v0.23.2) for other predictors, and compared the performance. Random forest performed comparably to other methods, and performed the best when tuned properly.

Negative samples were randomly downsampled to achieve 1:1 positive versus negative ratio.

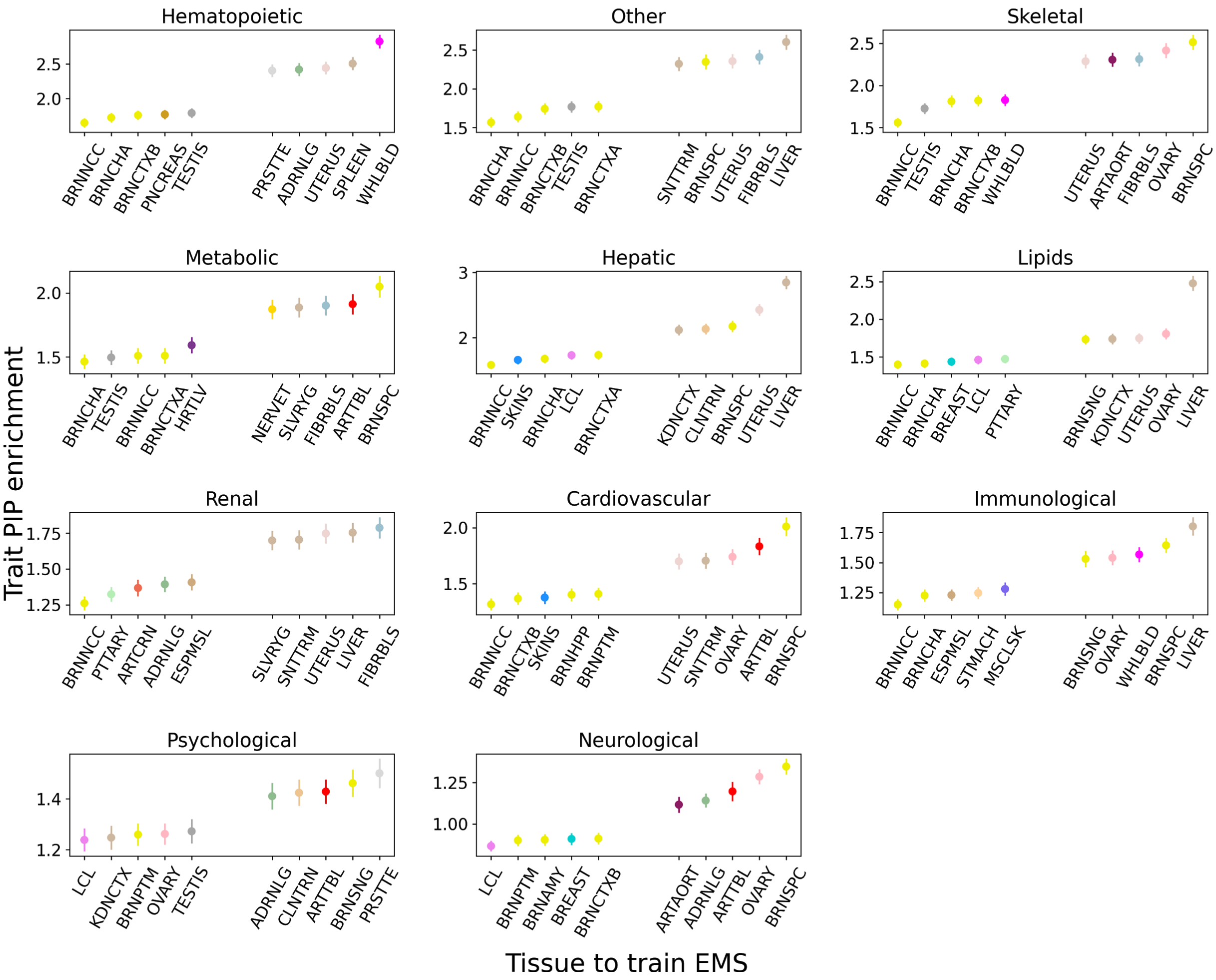


**Figure S21. Complex trait enrichments of top variant-gene pairs prioritized by EMS trained in different tissue**

Different panels correspond to different trait categories in UKBB. For each panel, the PIP enrichment in top 10,000 variant-gene pairs prioritized by EMS in the best and worst five tissues to train EMS are shown (complex trait PIP of each variant-gene pair is uniquely defined by the variant). Enrichment is compared to all the UKBB variants with PIP>0.0001, as in Ulirsch et al[1].
